# Expanding *N*-Glycopeptide Identifications by Fragmentation Prediction and Glycome Network Smoothing

**DOI:** 10.1101/2021.02.14.431154

**Authors:** Joshua Klein, Luis Carvalho, Joseph Zaia

## Abstract

Accurate glycopeptide identification in mass spectrometry-based glycoproteomics is a challenging problem at scale. Recent innovation has been made in increasing the scope and accuracy of glycopeptide identifications, with more precise uncertainty estimates for each part of the structure. We present a layered approach to glycopeptide fragmentation modeling that improves *N*-glycopeptide identification in samples without compromising identification quality, and a site-specific method to increase the depth of the glycoproteome confidently identifiable even further. We demonstrate our techniques on a pair of previously published datasets, showing the performance gains at each stage of optimization, as well as its flexibility in glycome definition and search space complexity. These techniques are provided in the open-source glycomics and glycoproteomics platform GlycReSoft available at https://github.com/mobiusklein/glycresoft.

## 1 Introduction

Protein glycosylation is the most heterogenous post-translational modification (PTM) [1, 2, 3] with effects on a wide array of biological processes [1]. Mass spectrometry has been established as one of the best tools for high throughput analysis of the glycoproteome [4]. Intact glycopeptide tandem mass spectrometry (MS/MS) interpretation is a challenging problem to address as data is generated with a variety of dissociation strategies and energies, and depending upon their appropriateness for different types of glycopeptides in glycoproteomes large and small [5, 6, 7, 8].

Depending upon the characteristics of liquid chromatography coupled tandem mass spectrometry (LC-MS/MS) used, different software strategies are needed [9]. As we and others have argued, high confidence glycopeptide identification requires good characterization and requires confident identification of the peptide and the glycan independently [10, 11, 12]. Stepped collision energy (SCE) collisional dissociation has been shown to be ideal for acquiring more complete fragmentation of *N*-glycosylated glycopeptides [8, 10, 5]. pGlyco2 [10] has been designed specifically with these characteristics in mind, including a glycopeptide spectrum match (GPSM) scoring model and multi-dimensional false discovery rate (FDR) estimation procedure for controlling the peptide and glycan FDRs jointly and independently. pGlyco2 uses a large database of glycan structures compiled from GlycomeDB [13] and other sources, to be able to exactly enumerate glycan fragments for a coverage calculation central to their method.

Collisionally dissociated glycopeptide MS/MS spectra are complex, having peptide b and y ions with or without glycan reducing end residues, glycan B ions, and intact peptide + glycan Y ions, with varying abundances in varying charge states depending upon the number and strength of the bonds in the precursor molecule and the collision energy used [6]. Thus far only glycan B ions have received critical analysis about their inter-relationships [14, 15] and limited work has been done on broad glycopeptide spectrum prediction [16] while these topics have been well explored and exploited in peptide spectrum matching [17, 18, 19, 20, 21, 22, 23].

We present a collection of methods to to learn inter-peak relationships based upon their fragment ion annotations, and to learn to predict the relative intensity of glycopeptide fragmentation events across a wide range of charge states. In addition, we present an generalization of our glycan network smoothing technique [24] to construct models of site-specific glycosylation to guide glycopeptide identification spanning the modeled sites. We provide an implementation of these techniques with GlycReSoft and its supporting libraries.

## 2 Methods

### 2.1 Datasets

We demonstrate our method on two previously published stepped energy HCD glycopeptide datasets. The first dataset, originally published by [10], was enriched glycopeptides from mouse brain (PXD005411), kidney (PXD005412), heart (PXD005413), liver (PXD005553), and lung (PXD005555) tissues, which we will refer to as the mouse tissue dataset. The second was originally published by [25] was enriched for sialic acid-containing glycopeptides from human serum (PXD005931) which we will refer to as the human serum dataset. The process we used involved a sequential refinement, but at each stage we used the same processed MS data, glycoproteome databases, and search parameters. The human serum dataset used a Thermo-Fisher Scientific Q Exactive with stepped NCE 15-25-35. The mouse tissue dataset used a Thermo-Fisher Scientific Orbitrap Fusion with stepped NCE 20-30-40.

#### 2.1.1 LC-MS/MS Preprocessing

We downloaded raw data files for each dataset from PRIDE [26], converted to mzML using ProteoWizard [27], followed by peak picking, deisotoping and charge state deconvolution using GlycReSoft’s preprocessing tool [28]. The preprocessing tool averaged each MS1 scan with the preceding and following MS1 scan, did not apply background reduction, used a glycopeptide averagine (H15.75 C10.93 S0.02 O6.47 N1.65) for MS1 scans and a peptide averagine for MSn scans.

#### 2.1.2 Database Construction

For the mouse tissue dataset, we used the UniProt Reference Mouse Proteome UP000000589 [29] and extracted only those from SwissProt. We extracted the *N*-glycan list from pGlyco2’s fixed database [10, 13] and simplified the entries from structures to compositions and combined it with a mammalian *N*-glycan biosynthetic simulation [30] allowing NeuAc, NeuGc, and Gal-*α*-Gal terminal groups. We combined this proteome and glycome, allowing one glycosylation per peptide, generating peptides using a trypsin cleavage rule allowing up to two missed cleavages, applied a constant carbamidomethyl modification at cysteine and variable oxidation modification at methionine. We generated decoy proteins by reversing complete protein sequences, but retaining *N*-glycosylation sites at the disrupted sequons without introducing new sites as in [10].

For the human serum dataset, we used the UniProt Reference Human Proteome UP000005640 [29] a human *N*-glycan biosynthetic simulation [30] allowing only NeuAc terminal groups, with a small curated list of *O*-glycans, but otherwise used the same as the mouse database.

#### 2.1.3 Search Strategy

For the mouse tissue dataset, we allowed a 10ppm mass error tolerance for precursor ion matches and 20ppm mass error tolerance for product ion matches, as done in the original study [10]. For the human serum dataset we used 10ppm mass error tolerance for both precursor and product ion matches and also allowed one ammonium adduct [28]. Our search strategy did not consider chimeric or co-isolating precursors, although when we compared results to pGlyco2, we included their chimeric solutions.

### 2.2 Base Scoring Model

We built upon the GPSM scoring model and FDR estimation paradigm developed in pGlyco2 [10]. The scoring model used a linear mixture of peptide backbone and glycan structure evidence to score glycopeptides. The peptide score (Eq. 1) was a mass accuracy weighted log-intensity summation, weighted by peptide sequence coverage (to exponent *γ*). The glycan score (Eq. 2) followed the same pattern, save that the glycan coverage is broken into two terms, where the coverage along the entire topology is given one exponential weight *α*, while the coverage of the conserved *N*-glycan core is given an additional exponent *β*. The two components are combined by a linear mixing weight *w.* Because the peptide and glycan scores were retained, the same mixture model-based FDR estimation procedure is applicable, allowing us to do a direct comparison with the results published in [10]. We used *α* = 0.5, *β* = 0.4, *γ* =1 and *w* = 0.65 for all variations of this scoring model.

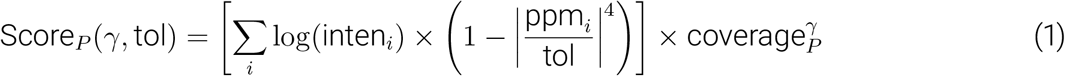

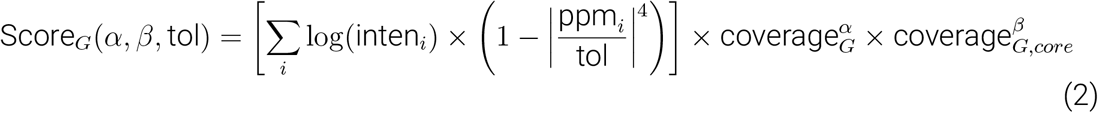

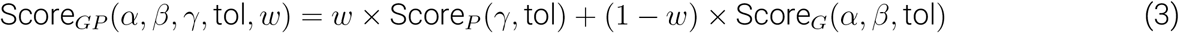

#### 2.2.1 Glycan Coverage Approximation

One advantage of pGlyco’s method is that it is able to compute a formal coverage ratio for the glycan component by using the peptide+Y ion ladder and an exact enumeration of the theoretical fragments for each of their glycan topologies. This comes at the cost of requiring a topology for each glycan to be searched, expanding the search space to consider, despite lacking diagnostic fragmentation to discriminate between most topological isomers. We introduce a method for approximating the total number of theoretical fragments a glycan composition could generate if its monosaccharides were arranged in a tree structure. *N*-glycans are branching structures, similar to binary trees. The height of a balanced binary tree with *n* nodes is log_2_ *n*. Because peptide+Y fragment generation often involves cleavage events along multiple branches, we can assume an upper bound for fragments of a binary tree to be *n* log_2_ *n*. *N*-glycans are not truely binary trees: the unfucosylated core motif’s root node has a single child node, suggesting the upper bound 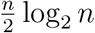 for *N*-glycans without core fucosylation or xylosylation. Beyond the first fan-out from the core motif, *N*-glycans are usually linear, causing the *n* log_2_ *n* approximation to be too harsh, especially for large glycans. A change to the natural log *n* log *n* turns out to be a close bound for small glycans and forgiving of large glycans which an exact coverage-based method is more stringent for. A comparison of the different rates of growth and divergence are shown in Figure 1. This allows us to generate a coverage metric for glycan compositions, letting us use a more compact glycan composition database rather than a glycan structure database.

**Figure 1:**
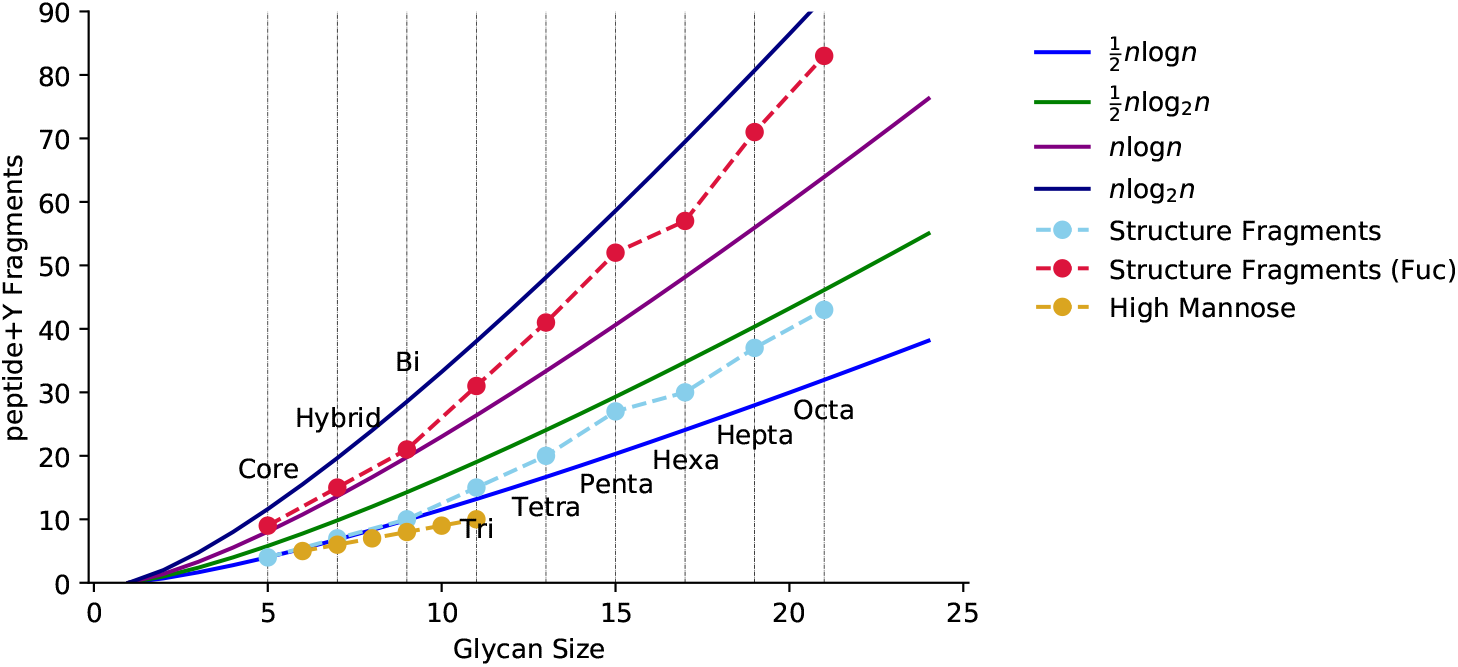
The number of distinct mass fragments produced by fragmenting a known topology of *N*-glycan of a given size, enclosed by our two approximation proposals, with and without the presence of core fucosylation. Each labeled vertical line is denotes a class of complex-type *N*-glycan of ascending number of lactosamine units beyond the core motif, which are arranged as separate branches. The immature high mannose from Man9 to Man4 are also shown, which fall below the approximation due to the homogeneity of their Y fragments.

We generated semi-structured peptide+Y ion ladders from glycan compositions by explicitly generating fragments assuming that a core motif is present, marking these fragments as *core* fragments, possibly with deoxyhexose or pentose side-chain, and then adding every combination of remaining non-labile monosaccharides to the core motif, including biosynthetically improbable ones. We treated NeuAc and NeuGc as labile. To calculate glycan coverage, we first approximated the “size” of a glycan composition as the number of monosaccharide residues in the glycan composition minus the number of NeuAc and NeuGc, and deduct one if two or more dHex/Fuc residues are present, calling the final number *n_g_*. We then use theüiycan bize approximation shown in Figure 1, computing the normalizing factor *d_g_* (Eq. 5).

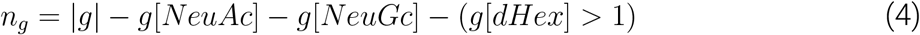

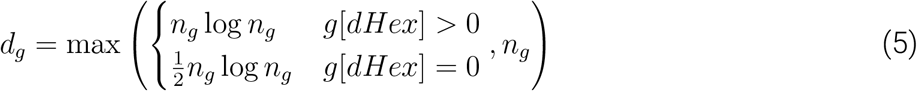

Where *g*[*m*] is the number of monosaccharide *m* units in *g* and |*g*| is the cardinality of *g*, the number of discrete monosaccharide residues in the glycan composition. The coverage_*G,core*_ is readily calculable as our algorithm explicitly enumerates the core fragments, while coverage_*G*_ is the number of distinct peptide+Y fragments matched divided by *d_g_*. Both target and decoy glycans were treated the same way, save that decoy glycans’ peptide+Y fragments beyond Y1 were given a random mass shift between 1.0 and 30.0 Da drawn from a uniform random distribution. We treated *O*-glycans identically, noting that for 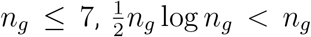, so by Eq. 5 *d_g_* = *n_g_*. Many individual mucin *O*-glycans are less than 7 monosaccharides. No glycans considered in our *O*-glycome were above this threshold.

#### 2.2.2 Supplementary Scores

We extend the aggregate score in Eq. 3 with two more features. The first is a mass accuracy bias (Eq. 6) with *μ_pre_* = 0 and *σ_pre_* = 5 × 10^-6^ to prefer solutions with better precursor mass matches, given equal fragmentation evidence, exploited in [28]. We add a penalty when a signature ion is expected for one of NeuAc or NeuGc but not observed, or when a signature ion for NeuAc or NeuGc is observed without being expected (Eq. 9) similar to [31]. Lastly, we include a small bias towards solutions with parsimonious localization in Eq. 10, preferring the solution with the best ratio of observed peptide b and y ions with a glycan fragment attached to the expected set given the position of the glycan where *n_p_* is the length of the peptide sequence. This new scoring function is expressed in Eq. 11.

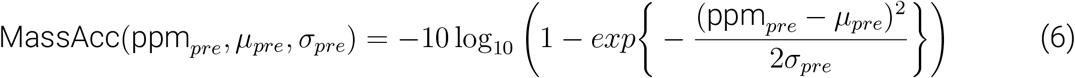

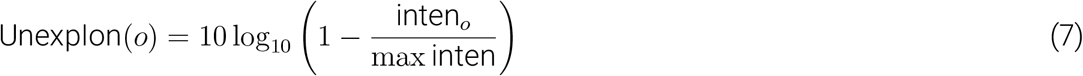

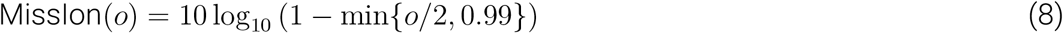

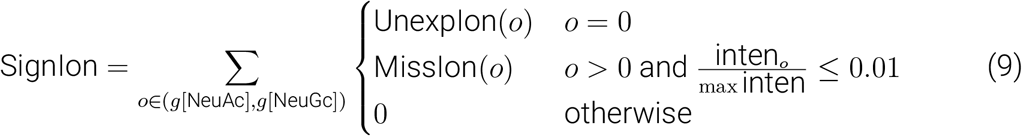

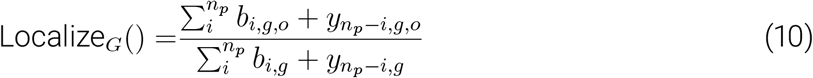

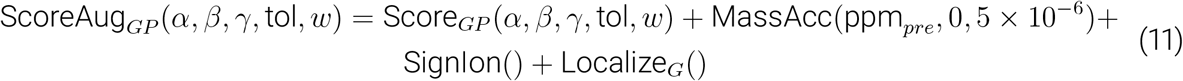

### 2.3 Inter-Peak Relationships

Glycopeptide fragmentation is complex, including multiple charge states for the same theoretical fragment and presence of both glycosylated and unglycosylated versions of peptide backbone fragments occurring interspersed. Many others [19, 17, 21] have demonstrated that a peak-fragment ion match which is supported by related peak-fragment ion matches is worth more than a peak matched in isolation. We used a Bayesian probability model based upon UniNovo’s [19]. In addition to the link features described in the original method, we added a mass difference feature for HexNAc (203.0794 Da) for peptide backbone fragments as well as for HexNAc, Hexose (162.0528 Da) and dHex (146.0579) for peptide+Y ion series matches. We did not include neutral losses of NH3 or H2O, complementary ions, and iterative refinement here, though they may be useful for future work. We set no restriction on peak rank as glycopeptide fragment ions are often low intensity.

UniNovo models multiple partitions over theoretical precursors independently by precursor mass as a proxy for peptide length assuming that fragmentation propensities for these structures will differ. As glycopeptides have both a peptide and a glycan component and a larger range of charge states than bare peptides, we use a multi-dimensional partitioning by peptide length, glycan size, precursor charge, precursor proton mobility [32], and the type and number of occupied glycosylation sites, with the ranges defined in Table 1. This produces up to 1500 partitions per glycosylation type, though not all are expected to be populated.

**Table 1:**
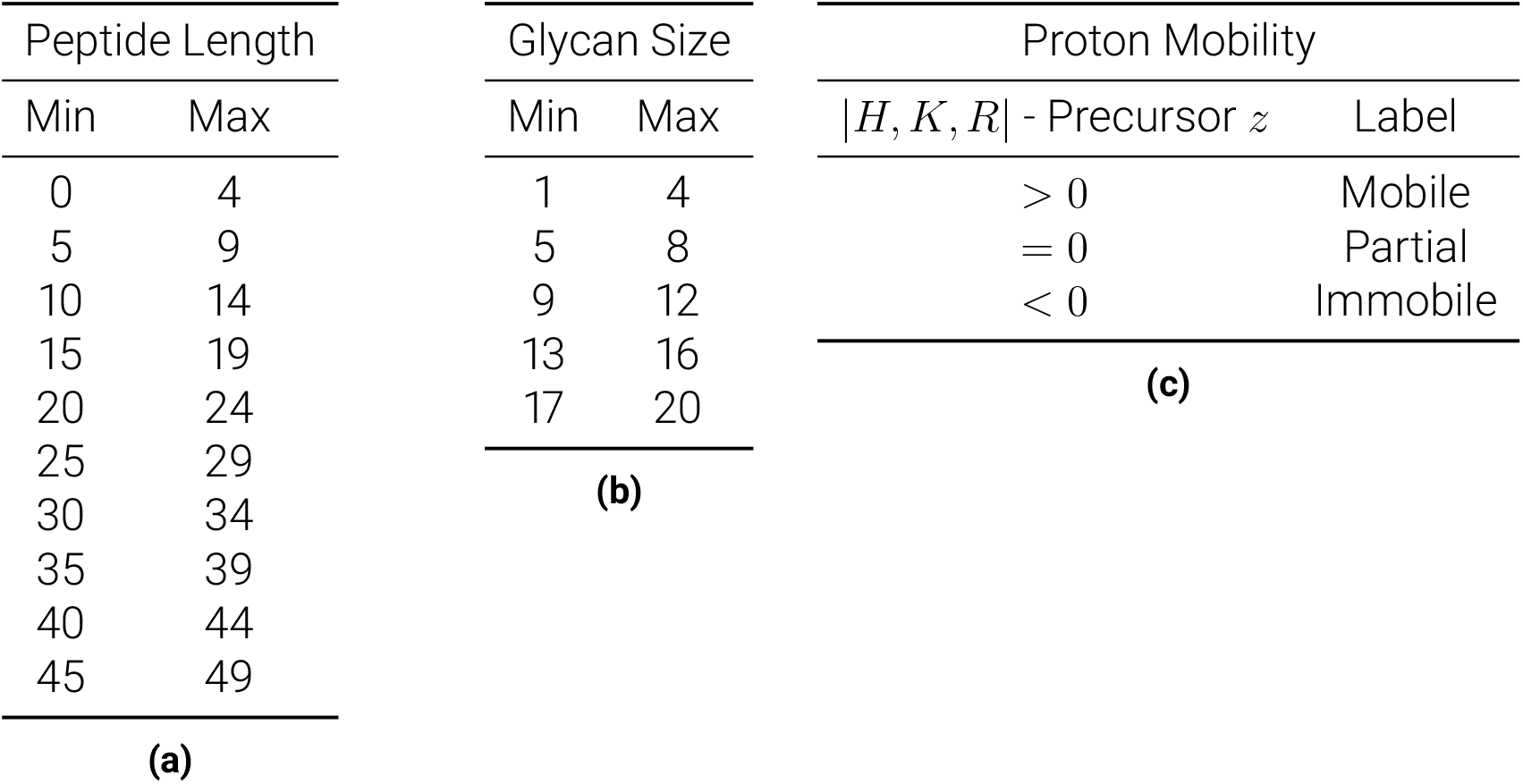
We partitioned glycopeptide spectrum matches from the training data into groups according to each of the ranges specified in Tables 1a,1b, 1c, their precursor charge *z* and the type and number of occupied glycosylation sites, producing up to 1500 theoretical partitions per glycosylation type.

We extracted GPSMs passing a 1% FDR threshold and a total score threshold of 20 from all samples in each dataset, converted to an annotated MGF format. For the mouse tissue dataset, we chose to reserve the brain tissue subset as a test set, and fit our model on the remaining tissue types, following the note in [10] that the glycopeptides found in each tissue type had small overlaps in the training tissues but no overlap with the brain tissue to demonstrate model performance and ability to generalize. For the human serum dataset there was no obvious distinction between samples, so we used all samples for training to demonstrate the effect of whole dataset modeling for a large study. We partitioned GPSMs according to the rules described above, though in order to smooth over small numbers of observations in some groups we mixed adjacent charge state groups while holding all other constraints constant, and fit our model for each ion series.

For each glycopeptide fragment match *f* we compute the posterior probability of that peak using its series and the set of unique peak-pair features which we term the reliability of the fragment match *ϕ_f_*.

### 2.4 Peak Intensity Prediction

A glycopeptide under collisional dissociation fragments in both the glycan and the peptide, with preference to weaker bonds breaking with greater frequency, subject to the physio-chemical properties of the molecule and the activation energy used [33, 34]. Prediction whether a fragmentation event should generate an abundant peak has repeatedly been used as a method for improving peptide identification [21, 32, 23, 22].

The mobile proton hypothesis is a widely accepted kinetic model of fragmentation for protonated peptides [33, 35] which has been used to create many peptide fragmentation prediction algorithms [21, 32]. Unsurprisingly, glycopeptide fragmentation depends on mobile protons as well, driving very different abundance patterns of fragmentation depending upon charge state [34, 6]. We used the proton mobility classification scheme described in [32], where the number of *K, R*, and *H* are compared to the precursor ion’s charge, where if the sum is greater than the charge, the precursor is *immobile*, if equal, the precursor is *partially mobile*, or less than, the precursor is *mobile.* We included this observation in the partitioning scheme we derived from UniNovo earlier, and apply the same partitioning scheme when modeling relative intensities.

We modeled the relative intensity of a fragmenting glycopeptide as a probability drawn from a multinomial distribution parameterized by a set of features listed in Table 2. The features chosen were based upon the approaches described in [21, 32]. It has been made clear that more sophisticated modeling techniques may be applied to this type of problem [22, 23], but they lack interpretability and require substantial numbers of observations to train, spectral appetites we cannot satisfy.

**Table 2:** We used these features to model the intensity of peaks associated with a glycopeptide fragmentation event. The “:” symbol denotes an interaction between two or more properties of a fragmentation event. The majority of these features are binary, with the exception of those starting with *stub glycopeptide:composition*, which may take on arbitrary non-negative integer values.

|  |  |  |  |
| --- | --- | --- | --- |
| n-term pro | stub glycopeptide:is_glycosylated 2 | stub glycopeptide:charge 1:glycan loss 4 | stub glycopeptide:charge 3:glycan loss 17 |
| n-term gly | stub glycopeptide:is_glycosylated 3 | stub glycopeptide:charge 1:glycan loss 5 | stub glycopeptide:charge 3:glycan loss 18 |
| n-term ser_thr | stub glycopeptide:is_glycosylated 4 | stub glycopeptide:charge 1:glycan loss 6 | stub glycopeptide:charge 3:glycan loss 19 |
| n-term leu_iso_val_ala | stub glycopeptide:is_glycosylated 5 | stub glycopeptide:charge 1:glycan loss 7 | stub glycopeptide:charge 3:glycan loss 20 |
| n-term asn | stub glycopeptide:is_glycosylated 6 | stub glycopeptide:charge 1:glycan loss 8 | stub glycopeptide:charge 4:glycan loss 0 |
| n-term his | stub glycopeptide:is_glycosylated 7 | stub glycopeptide:charge 1:glycan loss 9 | stub glycopeptide:charge 4:glycan loss 1 |
| n-term arg_lys | stub glycopeptide:is_glycosylated 8 | stub glycopeptide:charge 1:glycan loss 10 | stub glycopeptide:charge 4:glycan loss 2 |
| n-term x | stub glycopeptide:is_glycosylated 9 | stub glycopeptide:charge 1:glycan loss 11 | stub glycopeptide:charge 4:glycan loss 3 |
| c-term pro | stub glycopeptide:is_glycosylated 10 | stub glycopeptide:charge 1:glycan loss 12 | stub glycopeptide:charge 4:glycan loss 4 |
| c-term gly | stub glycopeptide:composition pro | stub glycopeptide:charge 1:glycan loss 13 | stub glycopeptide:charge 4:glycan loss 5 |
| c-term ser_thr | stub glycopeptide:composition gly | stub glycopeptide:charge 1:glycan loss 14 | stub glycopeptide:charge 4:glycan loss 6 |
| c-term leu_iso_val_ala | stub glycopeptide:composition ser_thr | stub glycopeptide:charge 1:glycan loss 15 | stub glycopeptide:charge 4:glycan loss 7 |
| c-term asn | stub glycopeptide:composition leu_iso_val_ala | stub glycopeptide:charge 1:glycan loss 16 | stub glycopeptide:charge 4:glycan loss 8 |
| c-term his | stub glycopeptide:composition asn | stub glycopeptide:charge 1:glycan loss 17 | stub glycopeptide:charge 4:glycan loss 9 |
| c-term arg_lys | stub glycopeptide:composition his | stub glycopeptide:charge 1:glycan loss 18 | stub glycopeptide:charge 4:glycan loss 10 |
| c-term x | stub glycopeptide:composition arg_lys | stub glycopeptide:charge 1:glycan loss 19 | stub glycopeptide:charge 4:glycan loss 11 |
| series b | stub glycopeptide:composition x | stub glycopeptide:charge 1:glycan loss 20 | stub glycopeptide:charge 4:glycan loss 12 |
| series y | stub glycopeptide:is_fucosylated 0 | stub glycopeptide:charge 2:glycan loss 0 | stub glycopeptide:charge 4:glycan loss 13 |
| series stub_glycopeptide | stub glycopeptide:is_fucosylated 1 | stub glycopeptide:charge 2:glycan loss 1 | stub glycopeptide:charge 4:glycan loss 14 |
| unassigned | n-term - 1 pro | stub glycopeptide:charge 2:glycan loss 2 | stub glycopeptide:charge 4:glycan loss 15 |
| charge 1 | n-term - 1 gly | stub glycopeptide:charge 2:glycan loss 3 | stub glycopeptide:charge 4:glycan loss 16 |
| charge 2 | n-term - 1 ser_thr | stub glycopeptide:charge 2:glycan loss 4 | stub glycopeptide:charge 4:glycan loss 17 |
| charge 3 | n-term - 1 leu_iso_val_ala | stub glycopeptide:charge 2:glycan loss 5 | stub glycopeptide:charge 4:glycan loss 18 |
| charge 4 | n-term - 1 asn | stub glycopeptide:charge 2:glycan loss 6 | stub glycopeptide:charge 4:glycan loss 19 |
| charge 5 | n-term - 1 his | stub glycopeptide:charge 2:glycan loss 7 | stub glycopeptide:charge 4:glycan loss 20 |
| series b:charge 1 | n-term - 1 arg_lys | stub glycopeptide:charge 2:glycan loss 8 | stub glycopeptide:charge 5:glycan loss 0 |
| series b:charge 2 | n-term - 1 x | stub glycopeptide:charge 2:glycan loss 9 | stub glycopeptide:charge 5:glycan loss 1 |
| series b:charge 3 | n-term - 2 pro | stub glycopeptide:charge 2:glycan loss 10 | stub glycopeptide:charge 5:glycan loss 2 |
| series b:charge 4 | n-term - 2 gly | stub glycopeptide:charge 2:glycan loss 11 | stub glycopeptide:charge 5:glycan loss 3 |
| series b:charge 5 | n-term - 2 ser_thr | stub glycopeptide:charge 2:glycan loss 12 | stub glycopeptide:charge 5:glycan loss 4 |
| series y:charge 1 | n-term - 2 leu_iso_val_ala | stub glycopeptide:charge 2:glycan loss 13 | stub glycopeptide:charge 5:glycan loss 5 |
| series y:charge 2 | n-term - 2 asn | stub glycopeptide:charge 2:glycan loss 14 | stub glycopeptide:charge 5:glycan loss 6 |
| series y:charge 3 | n-term - 2 his | stub glycopeptide:charge 2:glycan loss 15 | stub glycopeptide:charge 5:glycan loss 7 |
| series y:charge 4 | n-term - 2 arg_lys | stub glycopeptide:charge 2:glycan loss 16 | stub glycopeptide:charge 5:glycan loss 8 |
| series y:charge 5 | n-term - 2 x | stub glycopeptide:charge 2:glycan loss 17 | stub glycopeptide:charge 5:glycan loss 9 |
| series stub_glycopeptide:charge 1 | c-term + 1 pro | stub glycopeptide:charge 2:glycan loss 18 | stub glycopeptide:charge 5:glycan loss 10 |
| series stub_glycopeptide:charge 2 | c-term + 1 gly | stub glycopeptide:charge 2:glycan loss 19 | stub glycopeptide:charge 5:glycan loss 11 |
| series stub_glycopeptide:charge 3 | c-term + 1 ser_thr | stub glycopeptide:charge 2:glycan loss 20 | stub glycopeptide:charge 5:glycan loss 12 |
| series stub_glycopeptide:charge 4 | c-term + 1 leu_iso_val_ala | stub glycopeptide:charge 3:glycan loss 0 | stub glycopeptide:charge 5:glycan loss 13 |
| series stub_glycopeptide:charge 5 | c-term + 1 asn | stub glycopeptide:charge 3:glycan loss 1 | stub glycopeptide:charge 5:glycan loss 14 |
| is_glycosylated 0 | c-term + 1 his | stub glycopeptide:charge 3:glycan loss 2 | stub glycopeptide:charge 5:glycan loss 15 |
| is_glycosylated 1 | c-term + 1 arg_lys | stub glycopeptide:charge 3:glycan loss 3 | stub glycopeptide:charge 5:glycan loss 16 |
| is_glycosylated 2 | c-term + 1 x | stub glycopeptide:charge 3:glycan loss 4 | stub glycopeptide:charge 5:glycan loss 17 |
| charge 1:c-term pro | c-term + 2 pro | stub glycopeptide:charge 3:glycan loss 5 | stub glycopeptide:charge 5:glycan loss 18 |
| charge 2:c-term pro | c-term + 2 gly | stub glycopeptide:charge 3:glycan loss 6 | stub glycopeptide:charge 5:glycan loss 19 |
| charge 3:c-term pro | c-term + 2 ser_thr | stub glycopeptide:charge 3:glycan loss 7 | stub glycopeptide:charge 5:glycan loss 20 |
| charge 4:c-term pro | c-term + 2 leu_iso_val_ala | stub glycopeptide:charge 3:glycan loss 8 |  |
| charge 5:c-term pro | c-term + 2 asn | stub glycopeptide:charge 3:glycan loss 9 |  |
| series:c-term pro b | c-term + 2 his | stub glycopeptide:charge 3:glycan loss 10 |  |
| series:c-term pro y | c-term + 2 arg_lys | stub glycopeptide:charge 3:glycan loss 11 |  |
| is_glycosylated:c-term pro 0 | c-term + 2 x | stub glycopeptide:charge 3:glycan loss 12 |  |
| is_glycosylated:c-term pro 1 | stub glycopeptide:charge 1:glycan loss 0 | stub glycopeptide:charge 3:glycan loss 13 |  |
| is_glycosylated:c-term pro 2 | stub glycopeptide:charge 1:glycan loss 1 | stub glycopeptide:charge 3:glycan loss 14 |  |
| stub glycopeptide:is_glycosylated 0 | stub glycopeptide:charge 1:glycan loss 2 | stub glycopeptide:charge 3:glycan loss 15 |  |
| stub glycopeptide:is_glycosylated 1 | stub glycopeptide:charge 1:glycan loss 3 | stub glycopeptide:charge 3:glycan loss 16 |  |

For each partition of the training data, without allowing sharing between adjacent charge states, we estimated the parameters of the multinomial distribution from glycopeptide spectrum matches using iteratively reweighted least squares, weighted by the total signal in each spectrum, with the individual peaks weighted by the reliability ***ϕ*** using the peak relation model for that partition, or **1** if this lead to an unstable solution. Following regression model fitting, we calculated the Pearson correlation *ρ* between model predictions and each GPSM, and each case with a correlation below 0.5 was added to a “mismatch” set which we fit a second regression model. After this step, each partition contained a peak relation model *ϕ*, a relative intensity model, and an auxiliary relative intensity model denoted ***θ***.

### 2.5 Integrating Modeling into Scoring Functions

We incorporated peak relation-based reliability and peak intensity prediction into the glycopeptide scoring model’s two moiety-specific branches. For each theoretical GPSM, we found the appropriate partition’s models, or the nearest partition if it were missing, and computed a responsibility *π_θ_* for each model in the partition based on a weighted Pearson residual shown in Eqs. 12-14.

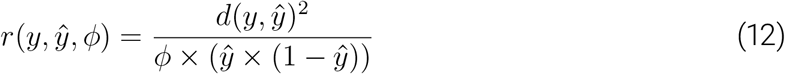

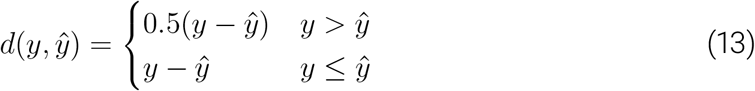

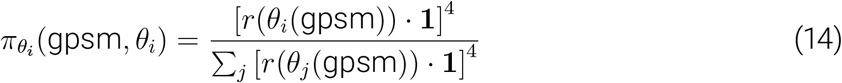

The prediction-enhanced scoring model extends the base scoring model with new components. The model peptide score shown in Eq. 17 gains three new components. We first transformed the Pearson residual *r* to a [0 - 1] scale using an empirically derived cumulative distribution function *R*, and convert this to an increasing logarithmic function in Eq. 16. The *e* function and *ϕ* channel weight away from improbable peaks, diminishing their influence on the score, while the correlation term gives a small benefit to matches which are larger while still consistent with the model prediction. We add a shifted Pearson correlation *ρ* of the observed and predicted intensities scaled by log_10_ *m_p_* where *m_p_* is the number of peptide fragments matched.

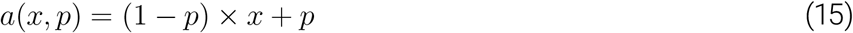

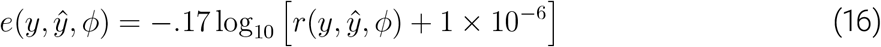

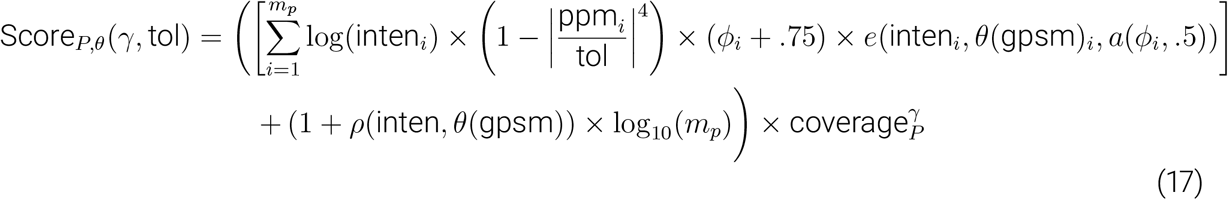

The model glycan score, shown in Eq. 18 is less changed to account for the much stronger relationship of model parameters with peptide+Y ions and to avoid putting too much weight on common but uninformative fragments. The 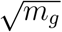 term being a compromise to be faster growing than log_10_ to accommodate for the lack of per-peak weighting, where *m_g_* is the number of peptide+Y ions matched. While the original scoring function in Eq. 3 had also included the `SignIon` term to prefer signature ion specificity, this addition to the glycan score now brings this to bear on the glycan FDR as well.

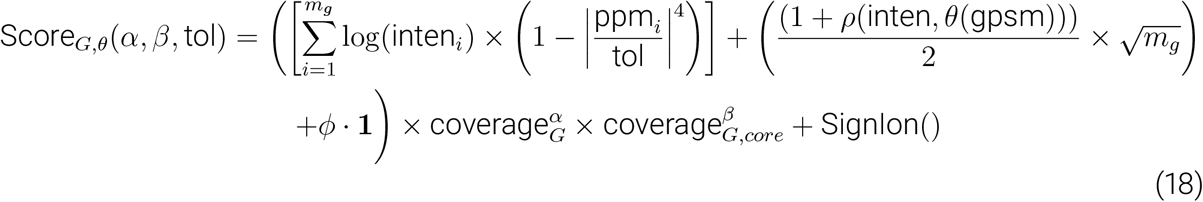

The we used the weighted sum of Eq. 17 for each relative intensity model *θ* for a partition, weighted by the mixture weights *π* for that GPSM to produce the peptide score. We do the same with Eq. 18 for the glycan score. The final scoring expressions are shown in Eqs. 19, 20, 21.

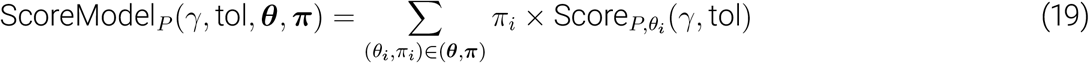

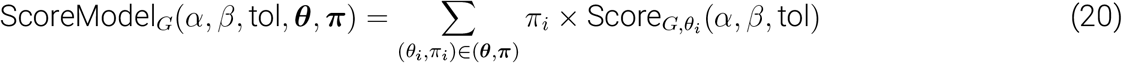

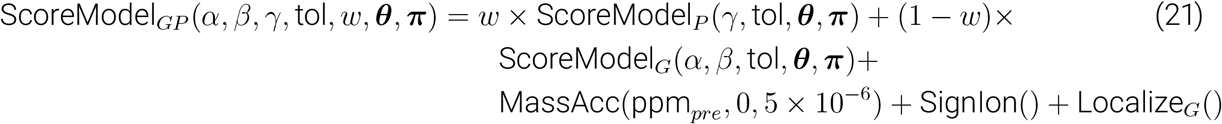

### 2.6 Site-Specific Glycome Network Smoothing

We and others have shown that it is useful to exploit the relationships between biosynthetically nearby glycans when evaluating glycan confidence [36, 24]. We extend the method applicable to released glycans we presented in [24] to the glycoforms observed at distinct sites of a glycoprotein, as determined by the high confidence identified glycopeptides spanning those sites, treating each site independently from all others. We generalize the approach to allow us to aggregate information across replicates by defining the variance of a glycan composition to be equal to 1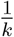 where *k* is the number of distinct times that glycan was observed at that site across peptides and across replicates. While the previous methods described learning a generalizable model of glycopeptide fragmentation, this method taps into the biological context of a sample. For the mouse tissue dataset, we fit site-specific glycome models for the brain tissue subset to be used with the test set for the fragmentation model scorer. For the human serum dataset, we again used the full dataset to estimate site-specific glycome models. The human serum dataset used the same *N*-glycan neighborhood rules found in [24], while the mouse brain tissue subset used an extended set of neighborhood bounds to take into account biosynthetic pathways absent in humans and other old world primates [1].

We computed the MS1 score *s* for each glycopeptide passing a joint FDR threshold of 1%, using the same features as in [24], except that we set the charge state score to a flat 0.8 prior to logit-transformation, as the charge state model was not appropriate for glycopeptides. While no MS2-based information is explicitly used to compute *s*, the 1% FDR threshold enforces that the glycan was observed confidently. We required each glycan be observed in at least two replicates for it to contribute to network smoothing parameter estimation. We used grid search to select the smoothing factor *λ* to estimate the network parameters *τ*, but computed the updated glycan confidence *ϕ_o_* and *ϕ_m_* using *λ* = 0.2 or the grid search *λ*, whichever was lower. Following model fitting, we saved a snapshot of the glycomes annotated with the network smoothed confidence values. To avoid notation ambiguity with the inter-peak reliability model, the symbol *ϕ* used in [24] to represent the smoothed glycan confidences, we denote the network smoothed glycan confidence **u** where **u**_*o*_ are the observed glycans that contributed to the model and **u**_*m*_ are the unobserved glycans informed by the model and **u**_*o*_.

To support using this method with decoy glycans, for each glycan in the glycome, we include a decoy glycan *g_id_* which has a distance of 1 to its analogous target *g_it_*, but a distance of 2 + *v* to all other target glycans where *v* is the distance between its analogous target *g_it_* and that other target glycan *g_v,t_*. Decoy glycans were otherwise treated identically, and participated in the neighborhood belongingness calculation and normalization and *u_m_* estimation.

To incorporate the site-specific glycome information into the search process, when we generate theoretical glycopeptides, we determine their protein and spanned glycosylation site(s) of origin, and look up the appropriate *u_g_* for that glycopeptide’s glycan *g* from that site’s model. Decoy glycans are looked up in the same manner, retrieving the decoy glycan’s *u_g_*. Decoy peptides are looked up against decoy proteins whose site models use the same site models reflected along the reversed sequence to align with the same residue, but otherwise behave identically.

We incorporated *u_g_* into the scoring function by adding it to the total evidence before being scaled by glycan coverage, as shown in Eq. 22, and use this as a replacement for Eq. 18 in Eq. 21, where *u_g_* = 0 if no site model was fit for that glycosite. We applied this modeling strategy to only *N*-glycosylation sites, no models were fit for *O*-glycosylation sites.

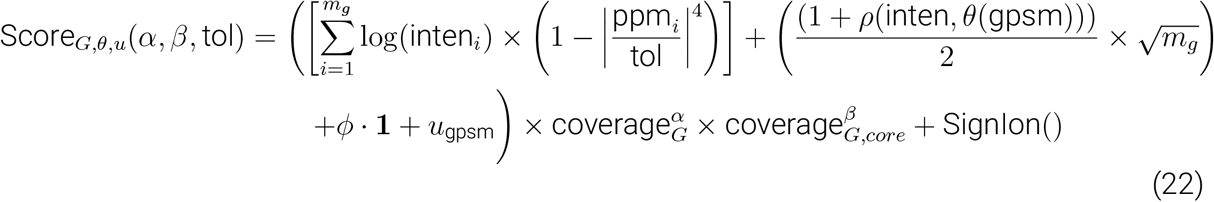

## 3 Results

### 3.1 Initial Search Results

The mouse tissue dataset produced 100,187 MS2 spectra passing the threshold required for model training, with 12,930 distinct glycopeptides, 3,119 distinct peptide backbones, and 599 distinct glycan compositions. The human serum dataset produced 39,797 spectra for model training, with 1,889 distinct glycopeptides, 664 distinct peptide backbones, and 118 distinct glycan compositions.

### 3.2 Glycopeptide ID Search Results

The number of GPSMs at 1% FDR for each algorithm run on the mouse brain dataset is shown in Figure 2. We also include the GPSM counts for these samples from [10] downloaded from PXD005411. The pGlyco2 results and our base algorithm are comparable, with pGlyco2 finding 97.6% of the GPSMS as our base algorithm. The search using our fragmentation modeling algorithm achieved a 21% improvement over the base algorithm, and the glycan network smoothing and fragmentation modeling algorithm yielded a 31% improvement over the base algorithm.

**Figure 2:**
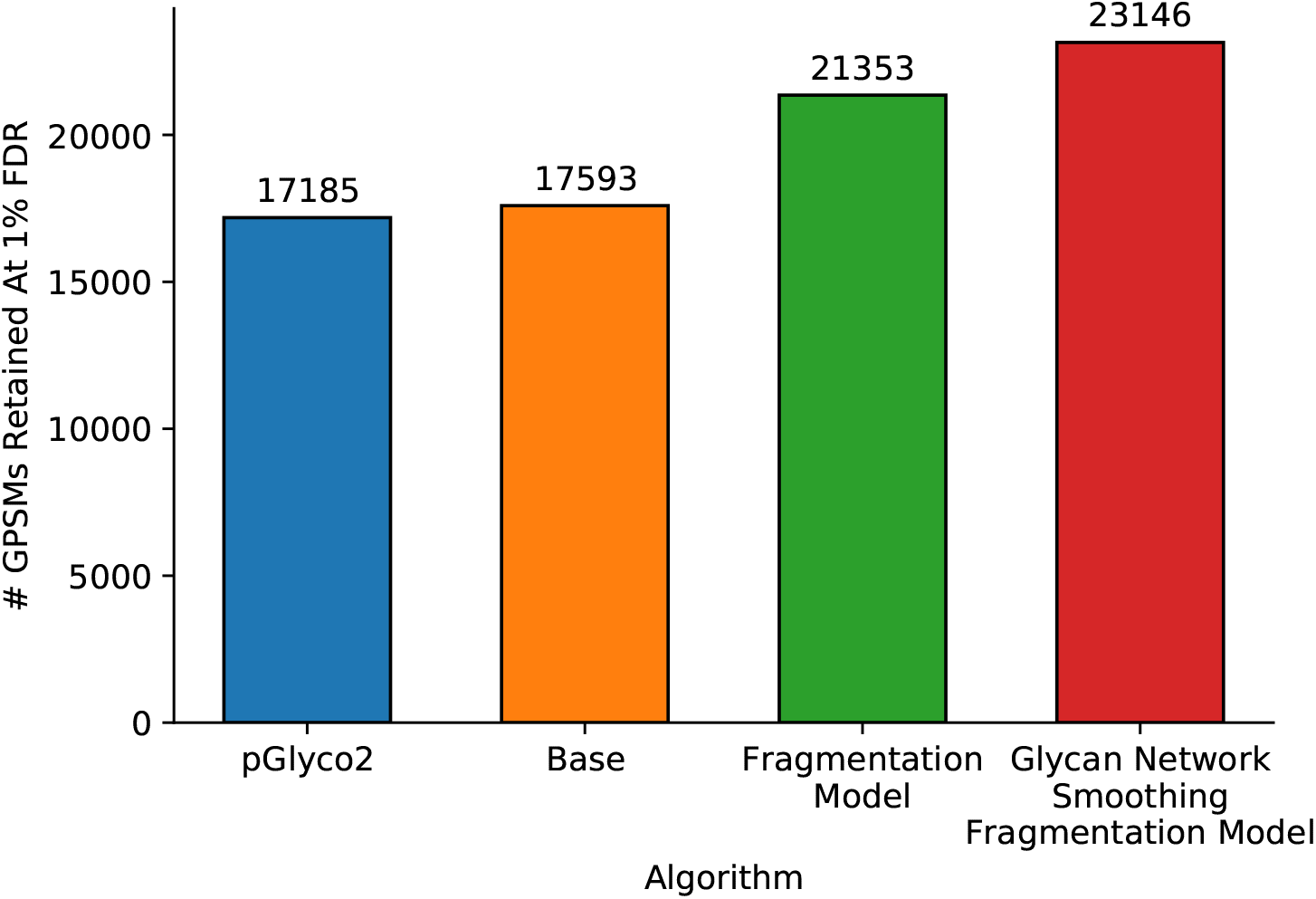
GPSM count at 1% FDR for each search algorithm from the mouse brain dataset. The pGlyco2 results from [10] and our base algorithm are comparable with pGlyco2 finding 97.6% of the GPSMS as our base algorithm, likely due to the expanded glycan database, while the search using our fragmentation modeling algorithm achieved a 21% improvement over the base algorithm, and the glycan network smoothing and fragmentation modeling algorithm yielded a 31% improvement over the base algorithm.

The number of GPSMs at 1% FDR for each algorithm run on the human serum dataset are shown in Figure 3. The fragmentation modeling algorithm yielded 16.1% more GPSMs over the base algorithm and the glycan network smoothing and fragmentation modeling algorithm achieved a 22.4% improvement over the base algorithm.

**Figure 3:**
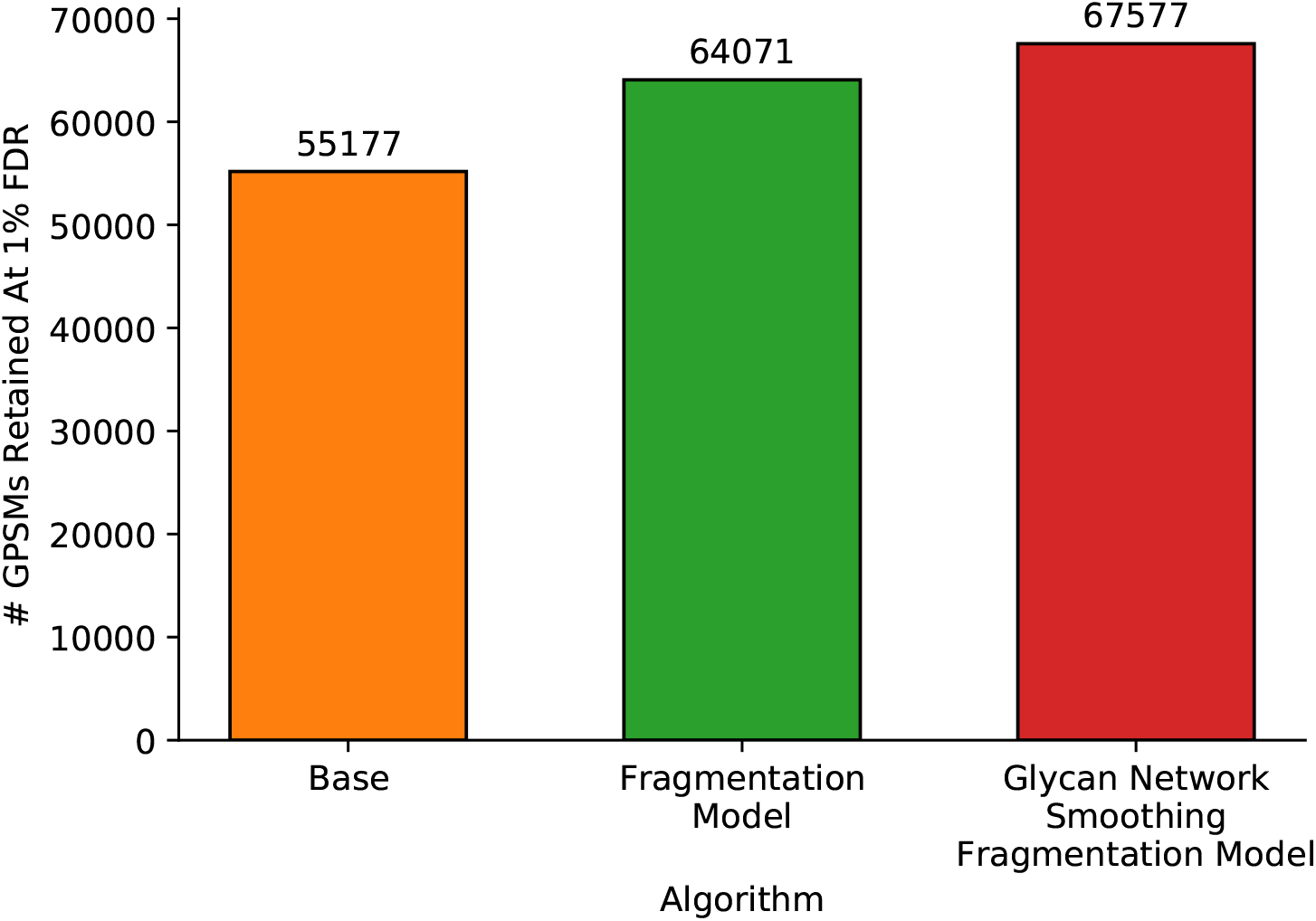
GPSM count at 1% FDR for each search algorithm from the human serum dataset. The fragmentation modeling algorithm yielded ?6.?% more GPSMS over the base algorithm and the glycan network smoothing and fragmentation modeling algorithm achieved a 22.4% improvement over the base algorithm.

### 3.3 Inter-Peak Relationships

The inter-peak fragmentation model for the mouse tissue model is shown in Figure 4a. The mean reliabilities for b, y and peptide+Y ion series were 0.23, 0.41, 0.95 respectively overall, and 0.18, 0.36, and 0.93 from the mouse brain subset data selected for modeling. The inter-peak fragmentation model for the human tissue model shown in Figure 4b. The mean reliabilities for these ion series were 0.23, 0.53 and 0.96 respectively.

**Figure 4:**
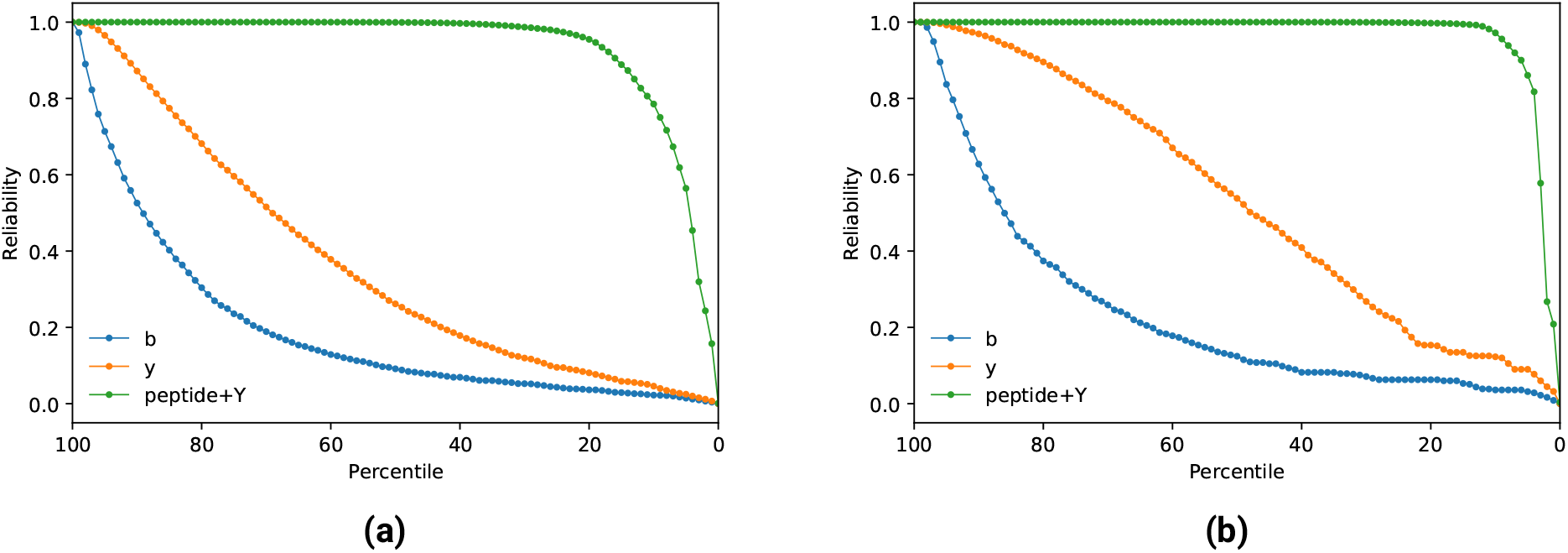
**4a** The trends of the reliabilities for the b, y, and peptide+Y ion series for the mouse tissue model drawn from the mouse brain subset selected for modeling. The mean reliabilities were 0.18, 0.36, and 0.93. 4b The trends of the reliabilities for the b, y, and peptide+Y ion series for the human serum model. The mean reliabilities were 0.23, 0.53 and 0.96.

### 3.4 Peak Intensity Prediction

The peak intensity model fit to the mouse tissue dataset, withholding the brain subset, predicted spectra with a median Pearson correlation *ρ* of 0.80 with their empirical spectra. When predicting on the brain subset alone, the median fell to 0.76. These relationships are shown in Figure 5a. The peak intensity model fit to the human serum dataset predicted spectra with a median correlation of 0.89 shown in Figure 5b. When considering the peptide b/y ions independently from the total correlation, the mouse tissue model has a median correlation of only 0.46, falling to 0.39 tested on the brain tissue subset. The peptide b and y ions make up a small fraction of the overall signal, but play a significant role in identification uncertainty estimation. The human serum model has a median correlation of 0.67 for the peptide b/y ions alone. Annotated spectra and their predictions are shown in Figure 6 showing examples of both common mobile proton examples but also partial and immobile proton examples (Figures 6d and 6c).

**Figure 5:**
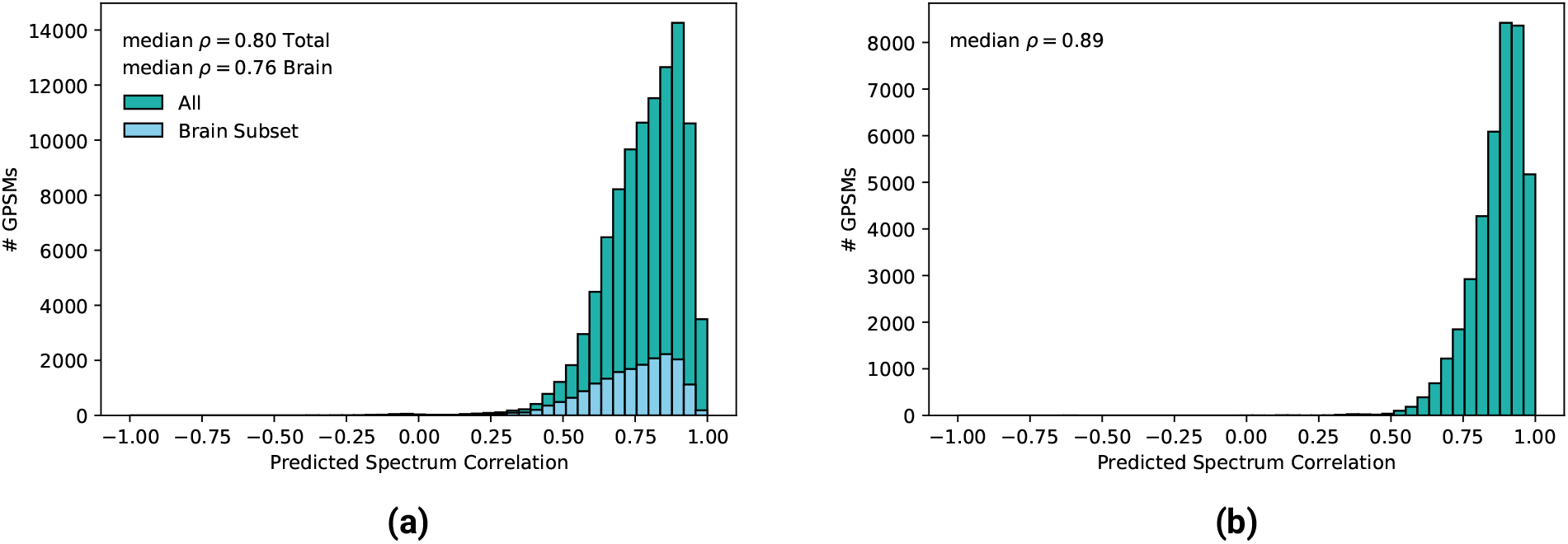
5a The distribution of predicted spectrum correlation of the mouse tissue median Pearson correlation over all data was 0.80, while only 0.76 on the brain tissue subset. 5b The distribution of predicted spectrum correlation for the human serum dataset with median Pearson correlation was 0.89.

**Figure 6:**
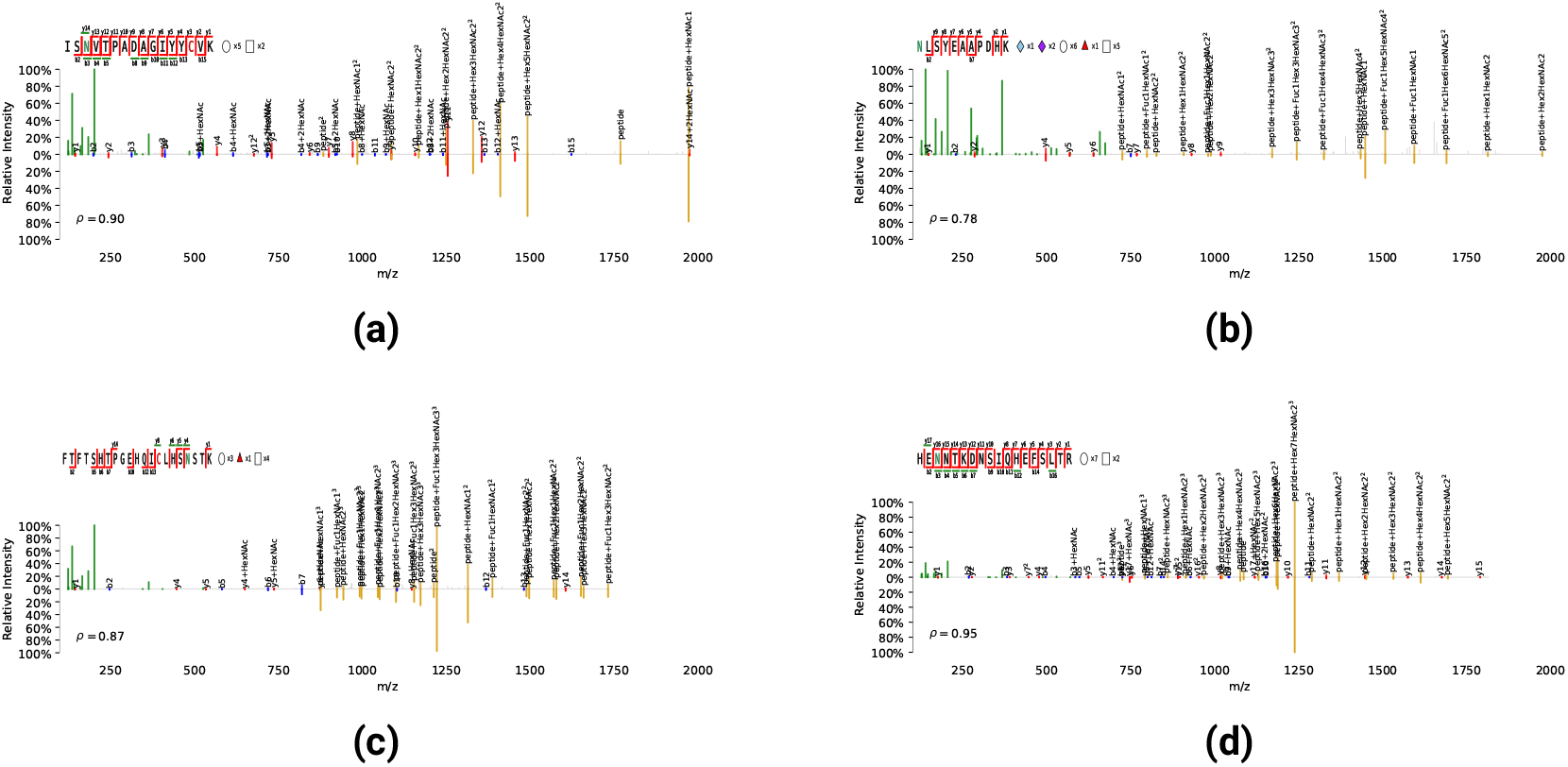
Annotated glycopeptides with their predicted spectra mirrored below them, and their Pearson correlation coefficient noted in the lower left, showing a range of glycopeptides. 6d shows an immobile proton match and 6c shows a partially mobile proton match, showcased by high charge state peptide+Y ions (yellow) and low abundance peptide b and y ions (blue and red). As we do not model oxonium ions (shown in green), these peaks are not included in the prediction

### 3.5 Site-Specifìc Glycome Network Smoothing

Using the site-specific glycan network smoothing model we estimated trends for 531 glycoproteins across 1075 glycosites observed in the mouse brain tissue dataset. We observed an emphasis for sites with *τ* > 1 for high mannose (337), hybrid (247), asialo-bi- (258), and asialo-triantennary (169) *N*-glycan groups, shown in Figure 7a. When applied to the human serum dataset, we modeled 223 glycoproteins across 356 glycosites. We observed an emphasis for sites with *τ* > 1 for bi-antennary (131), tri-antennary (129), hybrid (128), and asialo-bi-antennary (120), shown in Figure 7b.

**Figure 7:**
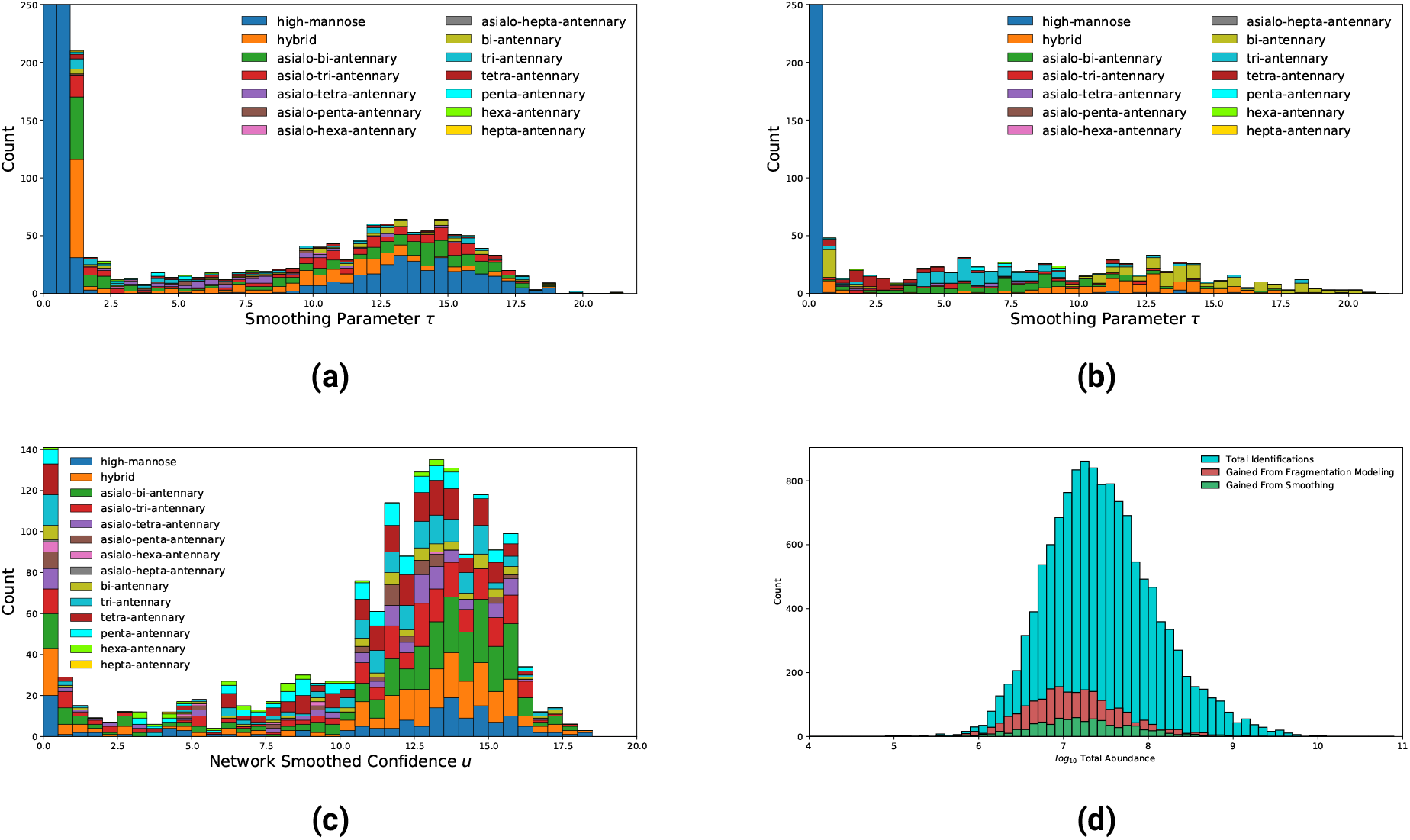
7a The distribution of network smoothing parameters *τ* estimated for the mouse brain tissue dataset as a stacked histogram. We observed an emphasis for high mannose, hybrid, asialo-bi- and asialo-tri-antennary *N*-glycans. 7b The distribution of network smoothing parameters *τ* estimated for the human serum dataset as a stacked histogram. We observed an emphasis for bi-antennary, triantennary, hybrid, and asialo-bi-antennary. The y-axis is truncated for both plots to show the non-zero site heterogeneity over the abundance of site-***τ*** pairs which were near zero.

The gained identifications from the addition of network smoothing to fragmentation modeling in the mouse brain dataset were assigned most often to the high mannose (154), hybrid (256), asialo-bi (311), asialo-tri (213), tetra (200), and tri-antennary (160). The network smoothing-gained identifications on average have a total abundance (4.5e7) of approximately half of the average total (1.02e8) for all identifications, shown in Figure 7d.

7c shows the neighborhoods that contributed to the identifications gained by network smoothing on the mouse brain tissue dataset as a stacked histogram. 7d shows the distribution of abundances for gained identifications from each stage of modeling overlaid atop each other.

## 4 Discussion

The work presented here demonstrates that it is possible to expand the number of high confidence glycopeptide identifications by a large margin without sacrificing stringency by incorporating physiochemical and biosynthetic factors into the identification process.

### 4.1 Inter-Peak Relationships

The inter-peak fragmentation modeling technique we derived from UniNovo [19] produced results similar to those observed in the original publication as shown by the reliabilities described in Figure 4, reaffirming the conclusion that peptide y ions are more reliable than b ions, with the strong but expected observation that peptide+Y ions tend to be extremely reliable, having consistent support from link features and charge difference features.

### 4.2 Peak Intensity Prediction

Our model’s ability to predict the relative intensity of individual product ions is comparable to previous rules-based methods [32], as shown in Figure 5, though its performance lags behind deep learning-based models for bare peptide prediction [23, 22]. While these more recent modeling approaches show much better performance, they were applied to a simpler problem and to hundreds of thousands to millions of training observations, while our training set is limited to less than 100,000 spectra per dataset.

The models from both datasets perform well on total glycopeptide fragmentation and on peptide+Y ion prediction, but the performance drops markedly for peptide b and y ions when considered in isolation. We used only 85 parameters to predict peptide b and y ions while we used 126 parameters to predict peptide+Y ions, though many of the peptide+Y parameters will be zero for most model partitions due to charge range interactions. We found no improvement in model performance attempting to use flanking amino acid features beyond the third position from the amide bond, while the gain going from the first to the second position away was modest at best. The amino acid context around a fragmentation site sets the stage for charge-local fragmentation pathway interactions [33, 35], which rule-based approaches to model fragmentation [21, 32, 37, 38] learned explicitly to a certain distance from the fragmentation site, while bi-direction recurrent neural network based methods [22, 23] learned across the entire sequence implicitly.

While the accuracy of the predictions vary, especially with respect to peptide backbone ions, in Figure 6 we show several annotated GPSMs with their predicted spectra mirrored beneath them. This also highlights the importance of the proton mobility feature. In low proton mobility settings, larger peptide+Y ions tended to be much more abundant, and all peptide+Y ions were present at higher charge states compared to high proton mobility cases, consistent with previous work [34, 6]. Our model seemed to perform well on the high mannose *N*-glycans of the mouse brain tissue dataset, but performed less consistently on sialylated glycans. This is likely because the model treated the loss of NeuAc/NeuGc as the same as loss of any other monosaccharide when these monosaccharides are much more labile.

The reason for the asymmetry between Eq. 17 and Eq. 18 was due to the overwhelming interaction between positively charged amino acid residues, proton mobility, and peptide+Y ions’ contribution to the score. A GPSM for a partially-mobile or immobile proton solution would over-emphasize the role that the peptide+Y ions played in the score, easily beating out a lower model concordance mobile proton solution that explained more of the experimental signal.

The identification performance gained from applying the trained model for glycopeptide identification yielded a 15 to 20% improvement displayed in Figures 2 and 3 in the number of GPSMs by penalizing solutions with improbable peaks at multiple levels, shrinking all scores but shrinking random match or decoy scores faster than targets without requiring a change to the FDR estimation procedure. The peak-level goodness-of-fit components play an important role in that shrinkage, while the flat additive features applied prior to coverage scaling help to add back some of what was lost when the entire match was higher quality.

The human serum dataset saw less benefit from the fragmentation modeling than the mouse brain dataset for two reasons. The primary reason was that the serum samples were simpler, having a sialic acid-enriched glycome and only a two hour gradient per sample, compared to the general glycome enrichment and four hour gradients used for the mouse samples. The second reason was that the human serum dataset was dominated by its glycan FDR, where unavoidable sharing between target glycans and decoy glycans of the often abundant peptide+Y0 and peptide+Y1 ions [7, 39] limited its influence. These ions are amongst the least specific, making them common in even poorly characterized glycopeptides. While the model also learned *O*-glycopeptides, there were relatively few distinct peptide sequences to learn from, so it is unlikely its performance is generalizable.

### 4.3 Site-Specific Glycome Network Smoothing

The site-specific trends we observed from the mouse brain tissue dataset were consistent with the observations by [10] and [4], showing few highly processed glycosites. The results from the human serum dataset were consistent as well [25, 40]. As previously published [24], the asialobi-antennary neighborhood allows up to one sialic acid due to the relationship with the hybrid neighborhood which does not discriminate between sialylation, and as such the structures would not be biosynthetically distant. The mammalian glycosylation network’s added terminal epitopes, NeuGc and Gal-*α*-Gal, played some role in how the more complex neighborhoods were estimated. Further, there may be undiagnosed adduction issues moving information from the sialylated glycoforms to the extra-hexosylated and fucosylated glycoforms.

This component contributes to gained identifications in two ways. The first is by spreading information between biosynthetically close-by glycan compositions reinforcing those glycans related to (and including) those in the training data, while the second works by decreasing the score at which a glycopeptide match would pass a particular FDR threshold by not raising the score of a truly random and unrelated matches. The first mechanism benefits only related glycans, though it can have its strongest effect on previously observed glycans. In the mouse brain tissue dataset the median networked smoothed observed glycan confidence *u_o_* was 13.6, while the median inferred network smoothed glycan confidence *u_m_* was 5.3 at 1% FDR. Assuming that false matches could be discarded by the 1% FDR threshold, multiple observation requirement, and the independent MS1 level score requirement may control for some but not all false matches, these may be reinforced by the smoothing method. However, glycan compositions that are supported by the network smoothing results are still required to show some glycan coverage to benefit from *u* (Eq. 22) and ***τ*** only benefits from multiple observations over multiple compositions, requiring systemic incorrect matches for the model parameters to be derailed substantially.

Our network smoothing glycan score (Eq. 22) at its core is using information external to the spectrum when scoring a GPSM, which violates the fundamental assumption of the targetdecoy approach [41]. We argue that the glycan coverage scaling factor reduces this as it forces matches to be increasingly non-random in order to benefit from external information, does not remove targets or decoys *a priori* from the search space, and is based on high confidence target matches which are then mirrored to each decoy type. Decoy glycans still participate in smoothing and the median difference between a target glycan and its decoy is 1.12, allowing a decoy glycan glycopeptide to out-compete its target counterpart if it matches another peak though ties, which contribute the majority of target-peptide-decoy-glycan matches, are no longer as common.

Although the human serum dataset included *O*-glycopeptides, we did not attempt to model mucin *O*-glycosylation here. The site-specific glycome network smoothing strategy is by its nature, specific to a site, which links it to localization. It can be difficult to reliably localize *O*-glycans with HCD alone [8]. A mucin *O*-glycan neighborhood definition would could be defined based upon related core motifs and extensions [2, 42]. This could make modification localization more biased, preferentially breaking larger *O*-glycans into smaller, previously observed *O*-glycans at different sites when no evidence is available for any configuration.

Site-specific glycan network smoothing is biologically derived, therefore more portable than fragmentation prediction, which is instrument and collision energy-dependent. It could be applied to other forms of glycopeptide identification tasks, such as the mixture spectra found in data independent acquisition datasets, provided there is some amount of peptide+Y fragmentation visible, or it could be used with another auxiliary metric of glycan evidence when peptide+Y ions are not present. Additionally, one could create a different front-end for the algorithm to consume information from curated glycome resources like GlyConnect [43], GlyCosmos [44] and GlyGen [45]. Such “literature-integrated” approaches would provide a rough approximation that a subsequent dataset-specific modeling round could benefit from.

### 4.4 Comparison With Reanalysis of Mouse Brain Tissue Dataset

As previously noted, the additional layers of modeling do not alter which product ions were searched for, only how their confidence is evaluated. Theoretically, these gains could be applied to different search methods as well, or connected with different FDR estimation strategies. Our method uses the multi-part FDR estimation strategy introduced by [10], which controls both peptide and glycan identification FDRs separately and jointly. Similar techniques have been proposed previously by combining multiple dissociation methods [12, 46]. Other techniques that use only a total score based upon a combination of evidence run the risk of identifying only one part of the molecule while leaving the other undefined, as [10] demonstrated with Byonic [31].

Recently, MSFragger-Glyco was published [47], which also re-analyzed the five mouse tissue datasets published in [10], that uses a total score FDR estimation strategy, but goes further to estimate the FDR for each glycan independently. MSFragger-Glyco reported over 40,000 GPSMs, more than double the number of glycopeptides identified compared to pGlyco2’s original results. This is explainable by using a total score based FDR rather than a multi-dimensional FDR. We extracted the number of GPSMs passing exclusively a 1% joint FDR, 1% peptide FDR, or 1% glycan FDR for [10]‘s and each layer of our search results, Figure 8, show that pGlyco2 yields 39,200 GPSMs, much closer to the number of GPSMs as MSFragger-Glyco reported. The number from our glycan network smoothing and fragmentation modeling result is even larger, exceeding 60,000. We argue that these identifications should not be considered high confidence because one of the moieties have not been identified.

**Figure 8:**
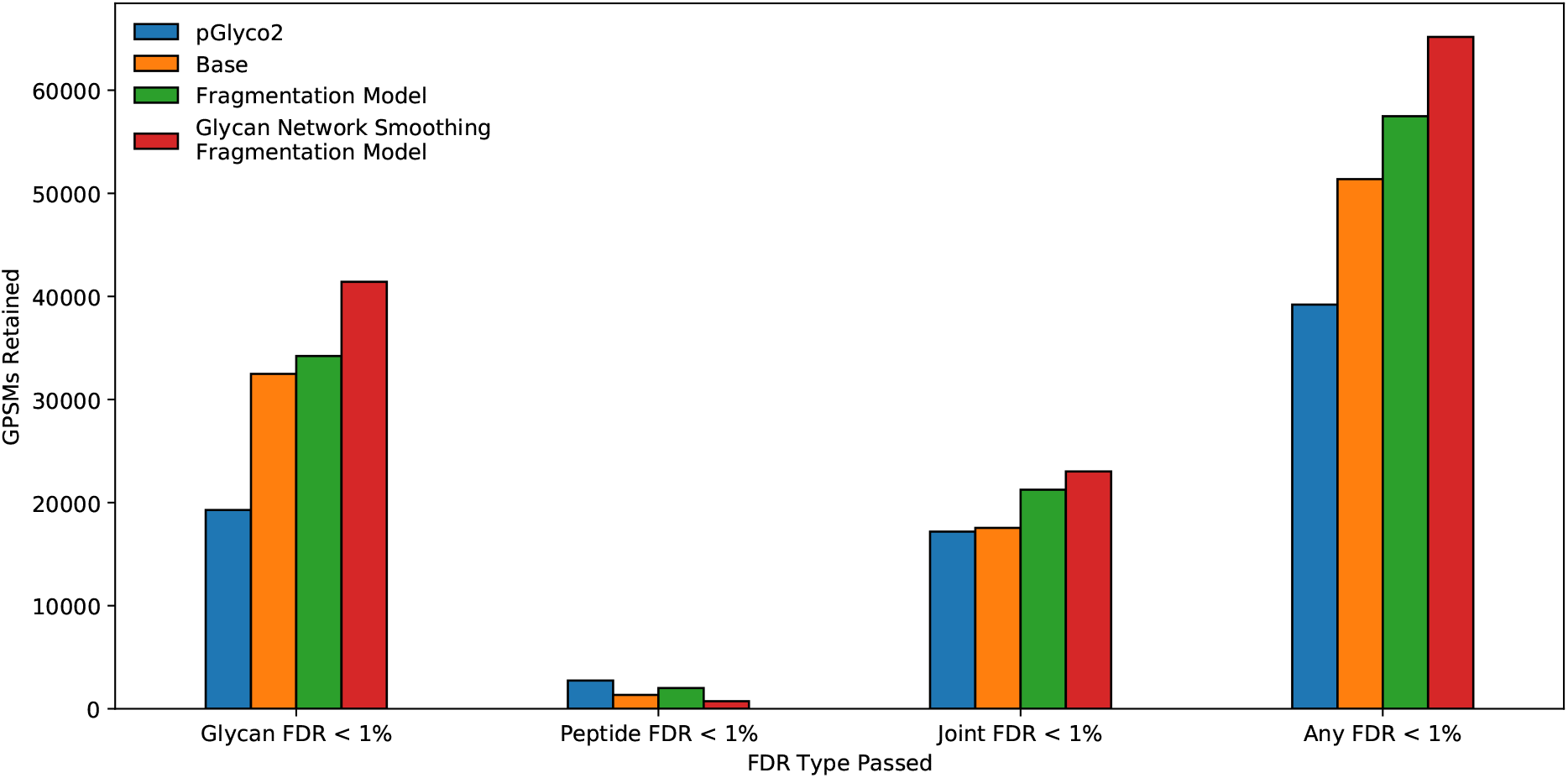
A comparison between the number of GPSMs retained exclusively by each FDR dimension at 1% FDR.

Consider an extreme match where a complete peptide+Y ion ladder is present, but we have not matched any peptide b/y ions. Such a spectrum match would score highly, but remains ambiguous across several peptide backbones. It is arguable that each distinct glycopeptide should be reported as tied matches for that spectrum and proceed, but such an identification is unhelpful for downstream analyses where we wish to quantify these precursors and make biological inferences. Alternatively, consider the case where a peptide backbone is fully characterized, but despite the fact that the glycan assigned with that peptide meets the precursor mass accuracy requirement, does not match any peptide+Y ions. Should that match be accepted, particularly if there are observable peptide+Y ion ladders in the spectrum not corresponding to that glycopeptide? There exist hypotheses that these types of identifications support, such as when analyzing a purified glycoprotein, but they do not represent identifications of the same quality as GPSMss which support both moieties.

MSFragger’s FDR model treats different mass shifts, proxies for any peptide modification, as distinct populations with different base rates of uncertainty [48]. This method work’s similarly to PeptideProphet’s ΔM component, combining a global expectation score dependent model with a mass shift-specific probability. While this approach is reasonable and for many classes of modification a necessity, the bin-specific probabilities may be learned from a small number of observations per mass shift. Supplementary table 8 of [47] shows that more than half of the mass shifts reported from the analysis of [4] were associated with less than 10 matches per mass shift. We observed a similar trend in our own identifications in the mouse brain tissue subset. We doubt that this has as large an impact on the number of identifications as the total score-based FDR estimate, as shown by Figure 8.

## 5 Conclusion

We present a method for improving glycopeptide identification by modeling their fragmentation behavior as well as the underlying biosynthetic process. We demonstrated an increased identification 15-30% yield at 1% FDR threshold depending upon sample complexity while maintaining stringency.

## 6 Code Availability

The source code for the search engine and network smoothing algorithm are part of GlycReSoft is available at https://github.com/mobiusklein/glycresoft. The fragmentation modeling codebase is available at https://github.com/mobiusklein/msms_feature_learning. Both codebases are released under the Apache 2 License.

## 7 Data Availability

The original datasets re-analyzed in this study are available from the PRIDE archive under the following accession numbers: mouse PXD005411 (brain), PXD005412 (kidney), PXD005413 (heart), PXD005553 (liver), and PXD005555 (lung) for [10] and from PXD005931 for [25].

