## Supplementary Figures for "Expanding *N*-Glycopeptide Identifications by Fragmentation Prediction and Glycome Network Smoothing"

### 1 Predicted Spectra

#### 1.1 *N*-Glycopeptide Spectra

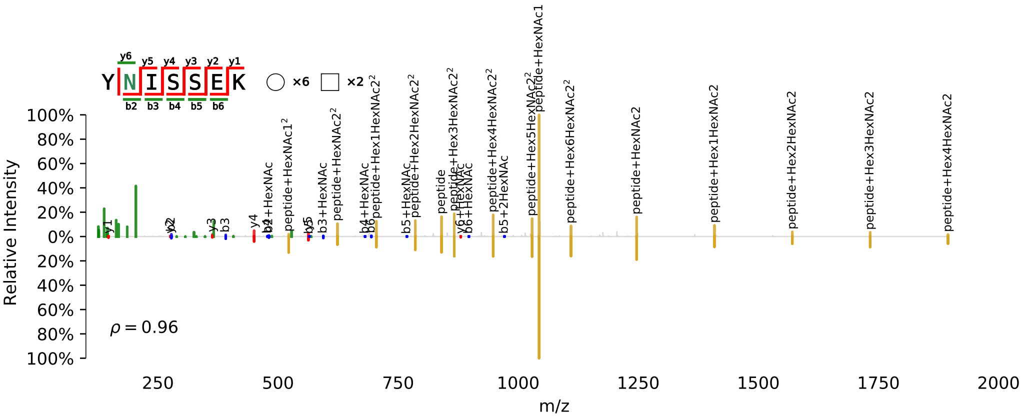


**(a)**

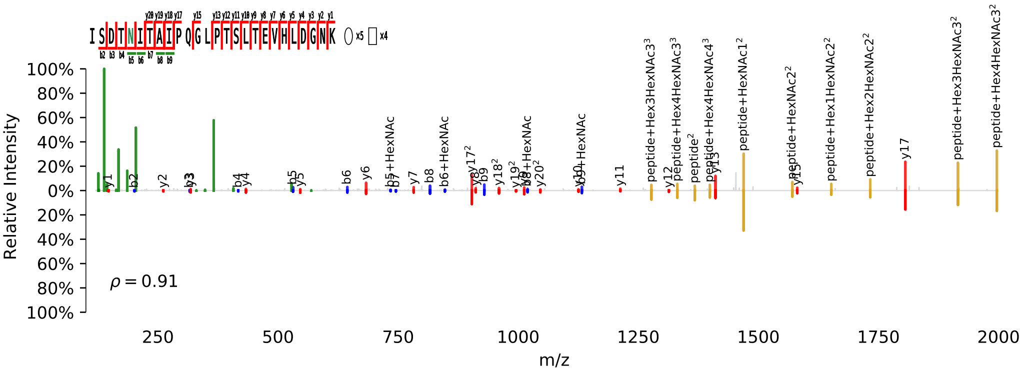


**(b)**

### 1.2 *O*-Glycopeptide Spectra

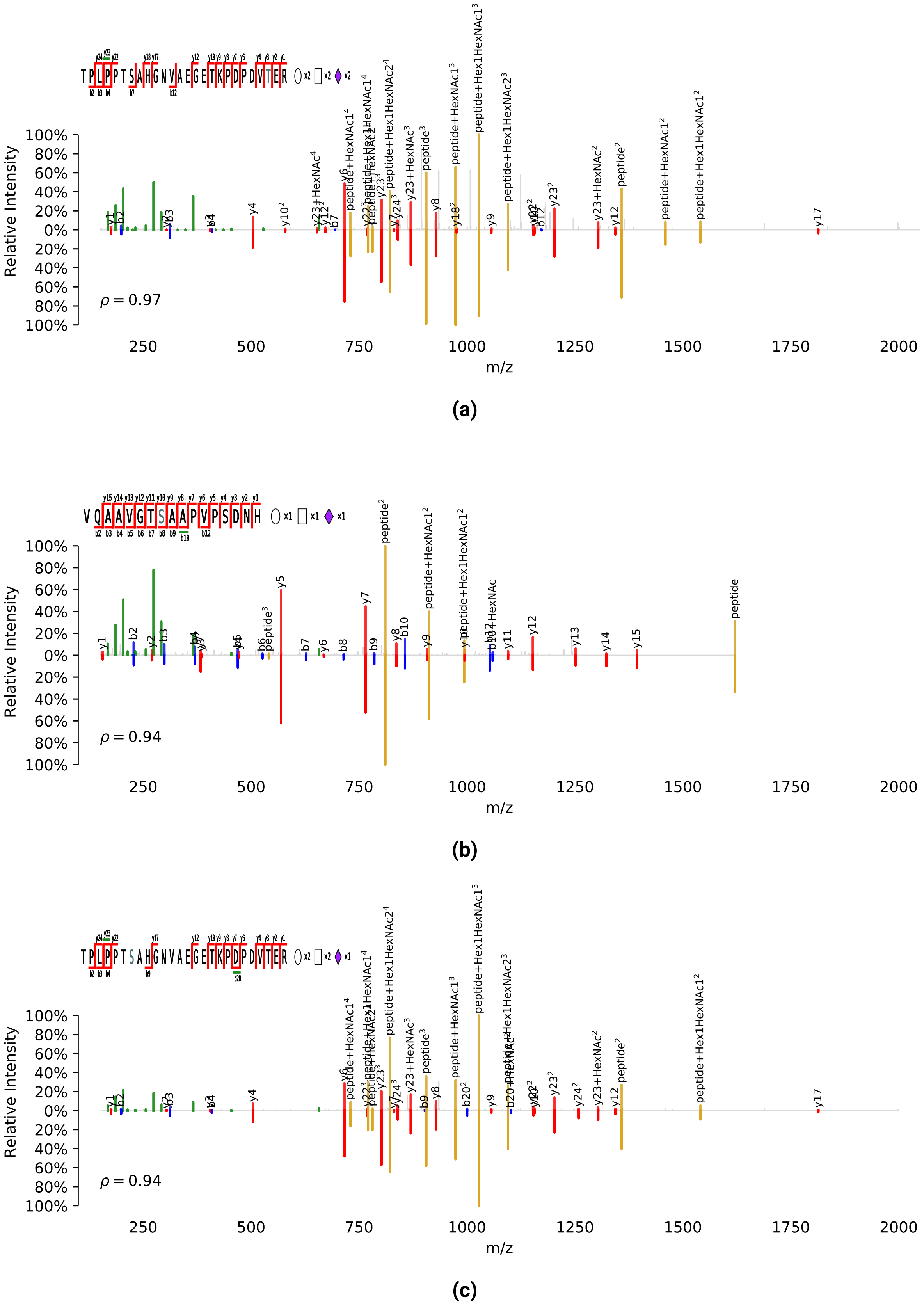


**Figure 2:** *O*-Glycopeptide spectral predictions appear strong, but they are likely over-fit due to the small amount of available training data for each partition
